# simExTargId: An R package for real-time LC-MS metabolomic data analysis, instrument failure/drift notification and MS2 target identification

**DOI:** 10.1101/151159

**Authors:** William M. B. Edmands, Stephen M. Rappaport

**Affiliations:** Rappaport Lab, UC Berkeley, School of Public Health, GL81 Koshland Hall, Berkeley, CA, 94720, USA.

## Abstract

The simExTargId R package provides real-time, autonomous, within-laboratory data analysis during a metabolomic LC-MS1-profiling experiment. Of concern to metabolomic investigators are instrumentation failure (especially for precious samples), outlier identification, instrument signal attenuation and pre-emptive feature identification for MS2 fragmentation.

SimExTargId allows observation of an experiment in progress with PCA plot and peak table outputs and also two shiny applications targetId for MS2 target identification and peakMonitor for signal attenuation monitoring. SimExTargId is ideally utilised on a (temporarily) dedicated workstation or server which is networked to a LC-MS data directory. Features include: email notification for instrument stoppage/drift, file format conversion, peak-picking, pre-processing, PCA-based outlier identification and statistical analysis. Additional MS1/MS2 experiments can be concatenated to a worklist or cleaning/recalibration undertaken if instrument drift is observed.

All source code and a vignette with example data are available on GitHub https://github.com/WMBEdmands/simExTargId/.

## Introduction

Collection of untargeted liquid chromatography-mass spectrometry (LC-MS) metabolomic datasets is subject to many potential pitfalls. For example, unobserved instrument failure can occur when an investigator is otherwise occupied or absent, this can be particularly poignant when samples are precious and/or limited. Subtle changes of instrumentation status may also escape notice such as slow leaks or partial blockages in LC systems leading to retention time drift and resolution loss. Column/ion-source build up or mass analyzer drift can lead to signal attenuation or resolution and mass accuracy loss. Other common concerns include pre-emptive outlier identification and statistically-relevant unknown MS^2^ fragmentation under the same instrumental conditions.

Recently, cloud-based approaches have emerged for autonomous data-processing however this necessitates upload of large data files to a remote server which also raises security and privacy concerns (Rinehart *et al.*, 2014; Warth *et al.*, 2017; Montenegro-Burke *et al.*, 2017). Scalable computational resources are now readily available for analysis of larger metabolomic datasets (e.g. 700+). For most analytical batches (i.e. 50-400 runs) even a relatively basic workstation (e.g. 64-bit Intel Core i7, 8GB Ram, 4 virtual cores) is able to tolerate repetitive peak-picking and data-analysis.

The R package *MetShot* facilitates nearly-online or “nearline” acquisition of MS^2^ spectra for features of statistical relevance (Neumann *et al.*, 2012). In contrast *simExTargId* provides a fully autonomous intra-laboratory workflow combining xcms and CAMERA followed by MetMSLine pre-processing, automated outlier detection and statistical analysis further decreasing the temporal gap from “nearline” to fully “online” MS^2^ data collection (Chambers *et al.*, 2012; Neumann *et al.*, 2012; Edmands *et al.*, 2014; Tautenhahn *et al.*, 2008; Kuhl *et al.*, 2012). An email notification mechanism also alerts users to instrumental stoppages or signal drift.

*SimExTargId* includes two web-applications, i.e. *peakMonitor* and *targetId* for signal monitoring and MS^2^ target list generation.

Mitigation for these pitfalls should be of broad interest to the metabolomic community.

## Results

An overview of the *simExTargId* autonomous workflow is shown in **Figure 1**.

**Fig. 1.**
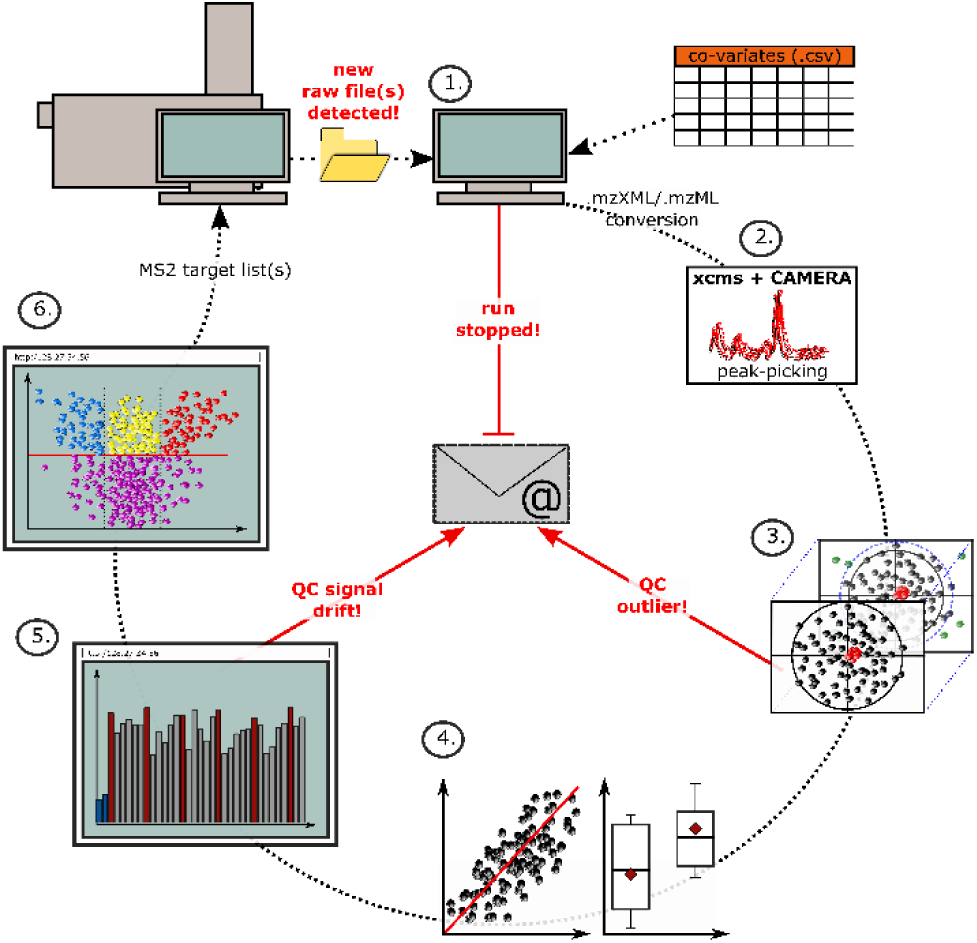
Overview of the autonomous simExTargId workflow and monitoring system.

### 2.1 Initiation, raw file detection and conversion

An example co-variate table (.csv) is provided with *simExTargId*. This worklist run order file inculcates an experimental design including column conditioning, MS^2^ data, blanks and pooled QCs. The first column is the file names and the second column the abbreviated experiment-types. Columns 3+ contain all co-variates to be considered for statistical analysis. A warning email is sent if the last data file was collected beyond a user-defined time (5 mins). Each raw data file is converted to the mzXML (or mzML) file format and saved in the auto-generated directories “/MS1” or “/MS2” (Chambers *et al.*, 2012).

### 2.2 xcms and CAMERA deconvolution

Peak-picking (xcmsSet) is performed file-by-file. After 10 samples retention-time correction, peak-grouping, peak-filling, pseudospectra and artefact annotation are conducted (Kuhl *et al.*, 2012). R workspace files (.RData) and peak tables are saved in sub-directories “R/rDataFiles” and “output/01.peakTables”, respectively.

### 2.3 MetMSLine pre-processing, PCA outlier detection and batch correction

The *MetMSLine* function *preProc* is used for pre-processing, with several steps being optional (Edmands *et al.*, 2014). Principal components analysis (PCA) is used to identify potential outliers (*pcaOutId*). If any of the outlying runs involve a QC sample an email is sent. PCA-based batch effect detection and adjustment is performed (*pcaClustId*). A weighted mean table for each CAMERA pseudospectra is also computed. Table and PCA-plot outputs are saved in the subdirectories “output/02.preProc” and “output/03.PCA” respectively.

### 2.4 Co-variate based automatic statistical analysis

Univariate statistical analysis methods are automatically selected for each co-variate (*coVarTypeStat*). Continuous variables and 2+ categorical variables are determined and then appropriate parametric or non-parametric statistical methods are applied. Output (.csv) is saved in the subdirectory “output/04.stats”.

### 2.5 Metabolite database peak-monitoring (application)

The function *peakMonitor* (optional) monitors previously identified metabolites supplied as a .csv file. The output can be also be viewed in a web-application. If a percentage (e.g. 80%) of the monitored-metabolite peak areas are found to be attenuated (e.g. −20%) then a warning email is sent.

### 2.6 MS^2^ target identification (application)

The application *targetId* visualizes the output in the directory “output/04.stats”, presenting volcano plots for all statistical analyses. Suitable samples for MS^2^ experiment are suggested based on peak area. Statistically relevant features for each co-variate can be downloaded and targeted MS^2^ experiment planned prior to MS^1^-profiling completion.

## Conclusions

*SimExTargId* is the first open-source package to provide comprehensive automation for metabolomic MS^1^-profiling data analysis. Peak-picking, data-deconvolution, pre-processing, PCA based outlier removal, batch detection/correction, statistical analysis and “online” MS^2^ target identification is performed in real-time and email notification provides system performance assurance.

